# Equations describing semi-confluent cell growth (II) Colony formation on a flat surface

**DOI:** 10.1101/2025.06.04.657949

**Authors:** Damien Hall

## Abstract

Individual cell growth can be affected by the presence of adjacent cells through a complex and multi-factorial biological process known alternatively as contact inhibition or confluence sensing. In a previous paper ***[Hall, D. 2023 Equations describing semi-confluent cell growth (I) Analytical approximations. Biophysical Chemistry, 307, 107173, pp. 1-9]***, sets of differential equations (with implicit analytical solutions) were developed to describe completely symmetrical cases of multicellular colony growth affected by variable levels of contact inhibition. Here we develop a model based on a spherical cap approximation of colony growth, that is able to describe variable contact inhibition for non-symmetrical multilayer cell formation on a solid plate. Although the model is realized as a set of interrelated ordinary differential equations, it is effectively governed by two parameters and is therefore capable for use in quantitative analysis of the kinetics of cell culture parameters such as shape, colony size and receding contact angle. The model is capable of accounting for transitions from monolayer to multilayer growth in a robust fashion.

## Introduction

The process of cell division results in the creation of a new cell directly adjacent to the original **[Su et al. 2012; Shao et al. 2017; Warren et al. 2019]**. If the newly formed cell is not subsequently moved to a new location (e.g. by diffusion, liquid shearing forces, or via innate cell motility) then the local environment of the parent cell may be altered by the presence of the daughter cell **[Adams, 1990]**. Such proximity effects caused by the presence of the new cell, on the growth behavior of the original cell, can be brought about by a myriad of different processes which include, local depletion of food resources **[Lavrentovich et al. 2013]**, restrictive mechanical effects **[Alessandri et al. 2013; Aland et al. 2015]**, secretion of biological regulatory factors **[Cooper et al. 1968; Hochberg and Folkman, 1972; Waters and Bassler, 2005]** and biochemical feedback circuits induced by the formation of cell-to-cell linker molecules **[Martz and Steinberg, 1972; Mendonsa et al. 2018, Ribatti, 2017]**. When cell-to-cell touching results in termination, or reduction, in the rates of cell growth and division, then the proximity phenomenon is known as contact inhibition (**Fig. 1a**) **[Abercrombie and Heaysman, 1954; Martz and Steinberg, 1972]**. The question of how cells respond both physically and biochemically to contact inhibition is fundamental to our understanding of research areas as diverse as the evolution of multicellular organisms **[Shapiro, 1998; Bassler and Losick, 2006; Ros-Rocher, 2021]**, bacterial biofilm growth **[Hartmann et al. 2019; Maier, 2021; Sauer et al. 2022; Pokhrel et al. 2024]**, optimization of cultured cell growth **[Kamath and Bungay, 1988; Galle et al. 2005]** and the study of cancer **[Huergo et al. 2012; Kapalczynska et al. 2016]**.

**Figure 1:**
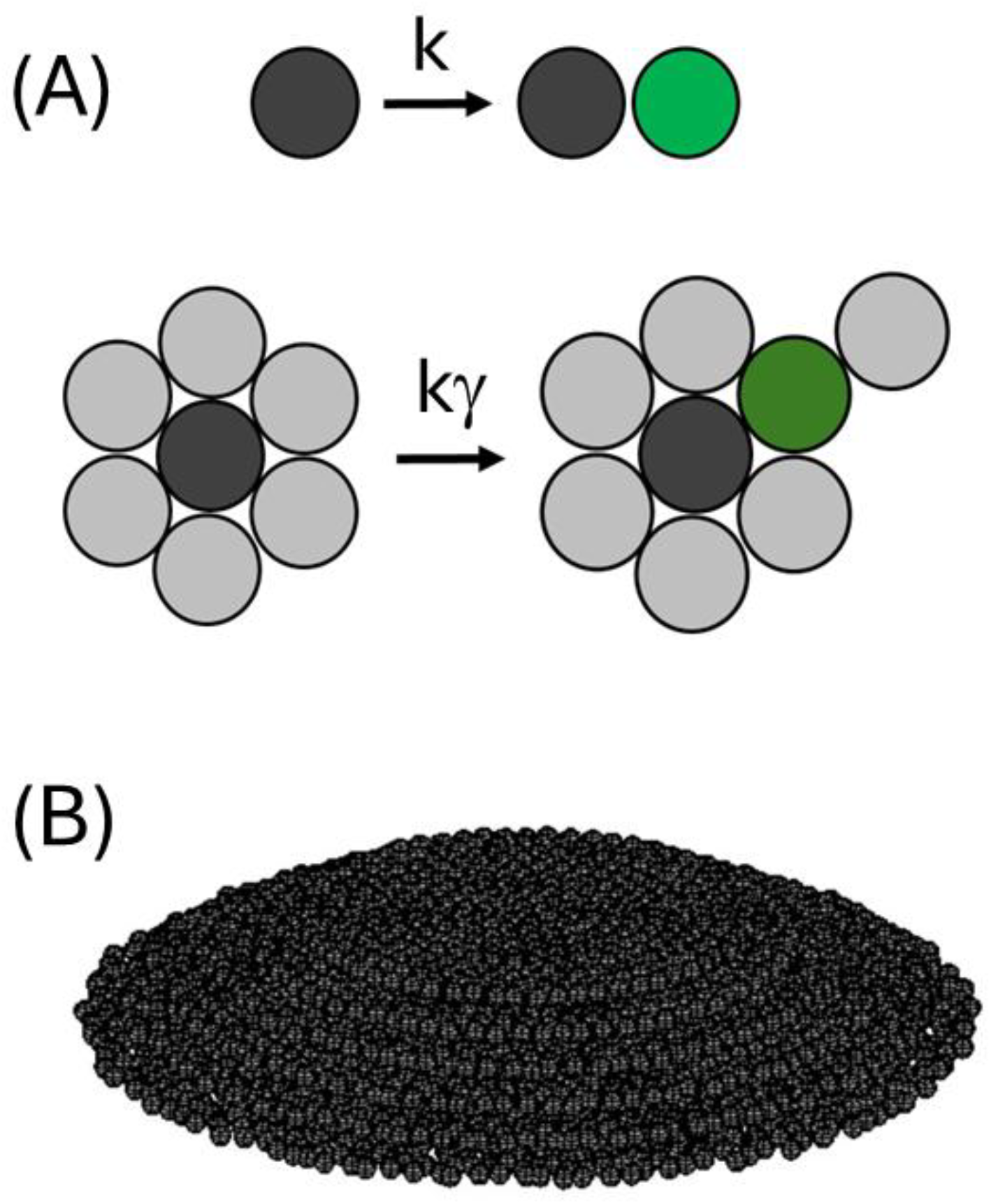
Effects of partial contact inhibition on cell division and growth **(A) Empirical lumped rate model of cell division/growth**: In the limit of infinite dilution, the rate of cell division and growth, in which one cell divides into two and the newly formed reaches maturity, can be described by a single lumped first order rate constant, k, that is ostensibly defined by intrinsic factors and bulk solution properties (upper). When surrounded by other cells, the rate of cell division and growth can be diminished by a factor γ (see Eqn. 2) due to a range of biochemical and physical effects collectively known as contact inhibition (lower). **(B) Schematic showing cell colony growth on a two-dimensional surface:** Partial contact inhibition of cell growth and division can result in the formation of a cell colony (or cell mass) having both area and height. Modelling the evolving shape properties of the growing colony is the subject of the present work.

In the previous paper in this series **[Hall, 2024]**, equations capable of describing reduced rates of cell growth and division for variable levels of contact inhibition were developed for the limited situation of symmetrical colony growth occurring in either two-dimensions (resulting in a circular monolayer), or three-dimensions (producing a spherical cell mass) **[Mayneord, 1932; Radszuweit et al. 2009; Montel et al. 2013; Hall, 2023; Hall 2024]**. The current paper treats the case of partial contact inhibited cell growth occurring on a hard surface – a situation which necessarily introduces an asymmetry to the colony growth along the vertical axis (**Fig. 1b**). In the following sections, we describe the development of the model before using it to simulate a number of relevant cases. We then discuss how closely the present simulations comport to experimental observations of cell colony growth **[Kamath and Bungay, 1988; Nguyen et al. 2004; Huergo et al. 2012; Pokhrel et al. 2024]**.

## Materials and Methods Section

Calculations described in the following theory section were carried out by use of original computer programs written in MATLAB R2024a. Programs are available upon email request to the author.

### Theory Section

As recognized nearly a century ago **[Monod, 1949; Hartwell and Unger, 1977]**, the rate of cell reproduction/growth under dilute growth conditions^1^ can be well described as a first-order process characterized by a lumped rate constant, k (units of s^−1^) (**Fig. 1a, Eqn. 1a**). Under such circumstances, the differential and integrated rate expressions describing the total number of cells over time, N(t), may be respectively written as **Eqn. 1b** and **Eqn. 1c**.

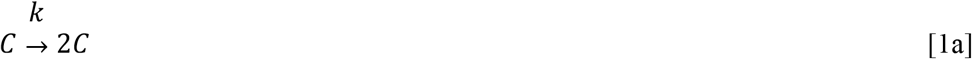

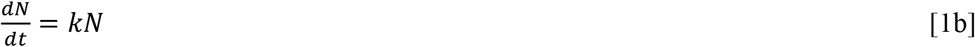

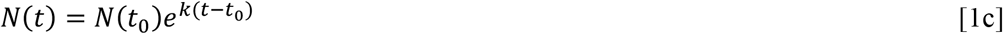

In previous work **[Hall, 2024]**, contact inhibition was defined by the immediate local density of cells around the cell of interest, with the extent of the effect set by the parameter β, such that β = 0 implies no contact inhibition and β = 1 dictates complete contact inhibition. To more closely align the mathematical notation with the physical process being modelled a reflective transition parameter, γ, was introduced such that a zero value of γ corresponded to a zero rate of cell division and a value of γ equal to 1 corresponded to unrestricted division and growth (**Eqn. 2a**) (**Fig. 1a**). In general, two subpopulations of cells were considered, the number of cells growing at the colony periphery, N_per_, and the number growing within the internal region of the colony, N_int_. Assignment of these subpopulations allowed for replacement of Eqn. 1b with **Eqn. 2b**.

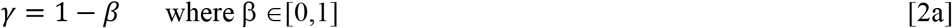

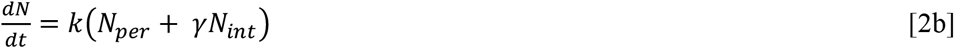

This approach allowed for straightforward simulation of cell growth subject to variable extents of contact inhibition for cases of completely symmetrical growth (i.e. for circular growth in two dimensions and spherical growth in three dimensions). However, it was not applicable to asymmetric growth patterns, such as those produced by cell colony growth on a hard surface, due to the fact that geometries of cell plated colonies do not adopt a simple fixed symmetry – meaning that colony size is not simply directly scalable with just the total cell number. In what follows we develop an argument, based on three assumptions, that allows for adaption of the prior treatment to simulation of cell colony growth on a hard surface.

#### Assumption 1: Directionality of the cell division/growth rate

For cells imbued with any intrinsic sense of directionality^2^ the components of cell division/growth rate from a single cell may be separately considered along the forward and reverse directions of each of the three Cartesian axes (**Fig. 2a**). From this perspective, the total cell division rate constant, k (as shown in Eqn. 1a) may be more properly expressed as a two-dimensional matrix consisting of six terms, **k**_**T**_ (**Eqn. 3a)**. For a single cell located on a culture plate, the hard surface will necessarily introduce a boundary condition to cell growth/ division along the vertical axis. However, in principle, no such boundary exists for growth/division parallel to the horizontal plane and as such, the problem of describing colony growth may be reduced to treating asymmetric growth along the vertical dimension. For these reasons it is convenient to further partition the rate constant matrix into two parts that relate to division and growth in the horizontal plane, **k**_**H**_ (**Eqn. 3b**) and the vertical plane, **k**_**V**_ (**Eqn. 3c**).

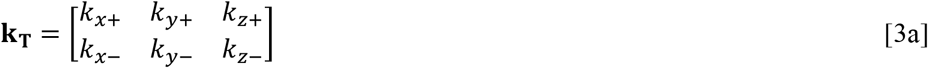

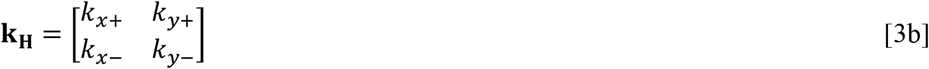

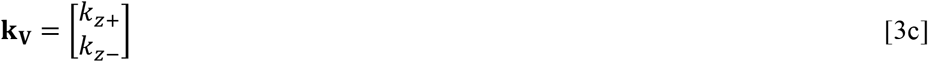

**Figure 2:**
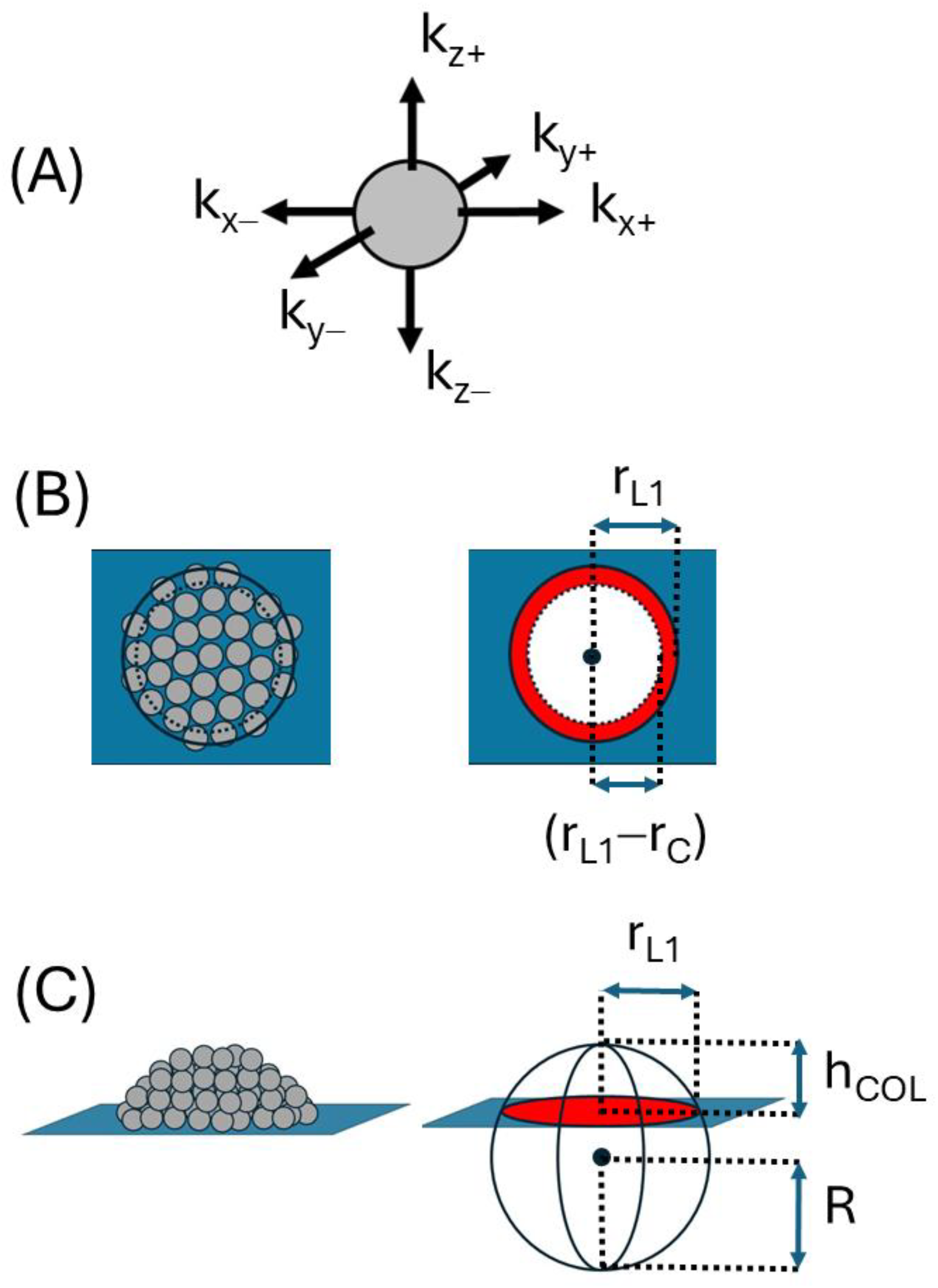
Three principal assumptions of the model **(A) Lumped rate constant, k, governing the cell division/growth process at the surface layer can be resolved into spatial vector components:** Rate constant can be expressed in matrix form (Eqn. 3) with components of these matrices grouped and summed to describe rate constants governing growth/division parallel, *k*_∥_, and perpendicular, *k*_⊥_, to the surface (Eqn. 4). Isotropic or anisotropic colony growth can result depending on the values of the individual matrix components. **(B) Growth of basal cell layer is circularly symmetric and occurs entirely to in plane growth:** Total number of cells in the basal layer, *N*_*L1*_, consists of two sub-types, cells at the perimeter, 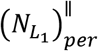, and cells growing internally, 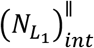, with the growth of the latter type subject to regulation by contact inhibition (Eqn. 5e). The basal cell layer is modelled as a circle of radius, r_L1_. Internal type cells are contained within the sub-circle of radius, r_L1_ − r_C_, while perimeter cells are contained within the circular segment existing between the radial domain [r_L1_ − r_C_, r_L1_], (Eqn. 5a-d). **(C) Cell colony grows as a spherical cap:** A spherical cap is the volume produced by intersection of a sphere and a plane, with the greater sphere defined by a radius R and the spherical cap defined by a circle of intersection, of radius r_L1_, and cap height above the intersecting plane, h_COL_.

Depending on the degree of directional anisotropy to the cell division/growth rate, each element of the rate constant matrix shown in Eqn. 3a can be assigned a fractional value, f, of the empirical rate constant, k, governing division of a single isolated cell indicated in Eqn. 1a. Assuming no intrinsic differences in growth/division rates along an individual axis (i.e. f_+_ = f_−_) and considering these fractions equal for the x and y axes (i.e. f_x_ = f_y_) then the aggregate rate constants describing cell division and growth parallel, *k*_∥_, and perpendicular, *k*_⊥_, to the surface plane, will respectively be given by **Eqn. 4a** and **Eqn. 4b**, with the relation between the fractional terms given by **Eqn. 4c**.

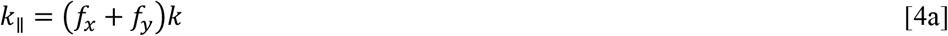

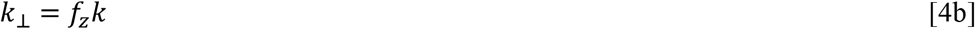

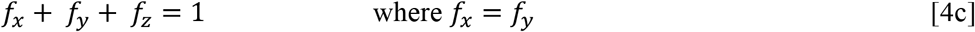

Within this schema isotropic growth is characterized by *k*_∥_/k = 2/3 and *k*_⊥_/k= 1/3, horizontally preferred anisotropic growth is defined by *k*_∥_/k > 2/3 and *k*_⊥_/k < 1/3, and vertically preferred anisotropic growth is defined by *k*_∥_/k < 2/3 and *k*_⊥_/k > 1/3.

#### Assumption 2: Base cell layer grows as a circle due to in plane growth

Lateral growth of the first layer of cells in contact with the plate is symmetrical and therefore will produce a circular shape, with the rate of this lateral growth governed by the rate constant, *k*_∥_, and the degree of contact inhibition, γ (**Eqn. 5**) (**Fig. 2b**) **[Hall, 2024]**. In practice, this assumption is equivalent to saying that there can be no increase in cells in layer 1 due to growth or division of cells in the layers above. We can relate the area properties of the colony base with the number of cells within it (**Eqn. 5a** and **5b**). In this formulation r_C_ is the radius of a single cell, 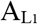 and 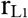 are respectively the area and radius of the bottom layer within the colony and F_2D_ is a unitless two-dimensional packing efficiency parameter that describes how much area each cell occupies in the circular area prescribed by the first colony layer. These geometric considerations allow for estimation of the number of cells contributing to cell growth and division along the horizontal plane of layer 1 located at the perimeter, 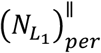, and internal, 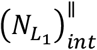 regions based on knowledge of the total number of cells in layer 1, 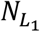, (**Eqn. 5c** and **5d**). These terms may be directly substituted into Eqn. 2b to produce **Eqn. 5e**.

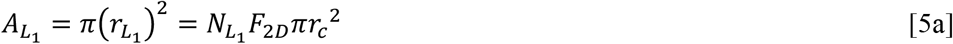

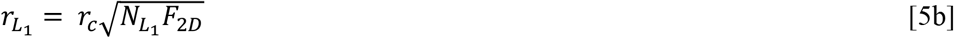

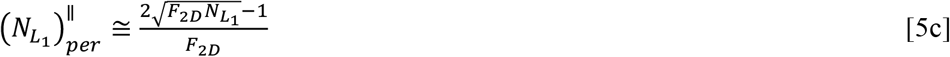

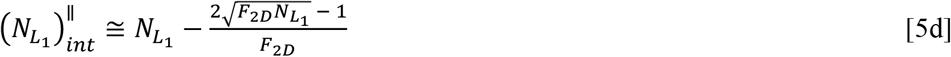

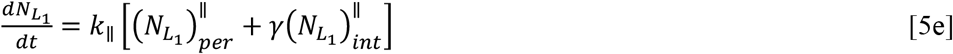

Numerical solution of Eqn. 5e yields 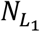 as a function of time from which its characteristic geometric parameters describing area and radius can be continually re-evaluated from Eqn. 5a and b.

#### Assumption 3: Colony grows as a spherical cap

At any stage the volume of the growing cell colony, *V*_*COL*_ (without reference to any shape consideration) is given by **Eqn. 6a** in which N_TOT_ is the total number of cells in the colony and F_3D_ is a unitless three-dimensional packing efficiency parameter that describes how much volume each cell occupies in the colony volume (**Eqn. 6b**).

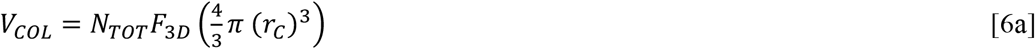

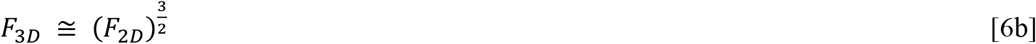

Due to the assumed equivalence of cell division/growth in all directions parallel to the plane, the growing plate colony is required to adopt a shape possessing lateral symmetry but not necessarily having vertical symmetry **[Palumbo et al. 1971; Kamath and Bungay, 1988; Nguyen et al. 2004; Galle et al. 2005; uergo et al. 2012; Su et al. 2012; Warren et al. 2019; Pokhrel et al. 2024]**. Amongst the smooth regular shapes this requirement is most simply satisfied by a spherical cap model i.e. the volume produced by intersection of a plane and sphere (**Fig. 2c**) **[e.g. see Palumbo et al. 1971; Kamath and Bungay, 1988; Nguyen et al. 2004; Pokhrel et al. 2024]**^3^. Aside from simplicity there is also strong experimental support suggesting that a large number of different bacterial and yeast colonies grow as a spherical cap for significant periods of their growth **[Nguyen et al. 2004; Pokhrel et al. 2024]** and that this shape approximation is also applicable to immortalized eukaryotic cell lines growing beyond confluence i.e. featuring anchorage independent growth **[e.g. see Lu et al. 1995; Galle et al. 2005; Nardone et al. 2011]**. In proceeding with this shape approximation we note that the circle of intersection of the spherical cap is effectively defined by the base cell layer, and as such the radius and area of the cap section are respectively given by 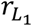 and 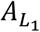 (**Eqn. 5**). The height of the spherical cap, relative to the growth surface, is defined as *h*_*COL*_. Using a cylindrical coordinate approach the volume of the spherical cap, *V*_*COL*_, can be expressed using Eqn. 7a (depending on the relation between *h*_*COL*_ and *r*_*L1*_) with Eqn. 7b describing the radius of the characteristic greater sphere, R.

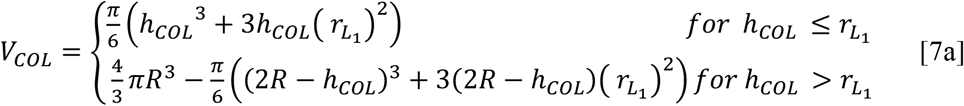

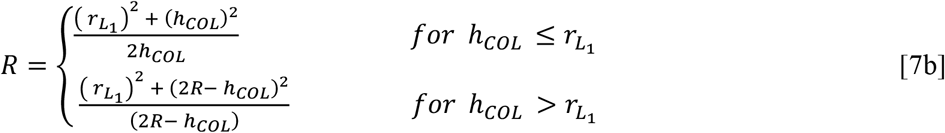

With the colony circular base radius and colony volume respectively given by Eqn. 5b and Eqn. 6a, Eqn. 7a can be solved iteratively to determine colony height, *h*_*COL*_, which can, in turn, be substituted into Eqn. 7b to return a value for the radius of the greater sphere, *R*. After calculation of the spherical cap volume it is then partitioned into two sub volumes (**Eqn. 8a**), the basal cell layer volume (**Eqn. 8b**) and the upper spherical cap volume, *V*_*UC*_, calculated by applying the functional form of Eqn. 7 with a reduced colony height, *h*_*COL*_*’* (where 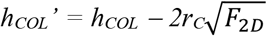) and a reduced radius of the circle of intersection, 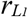 (where 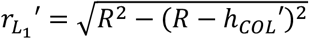) (**Eqn. 8c**) (**Fig. 3a**).

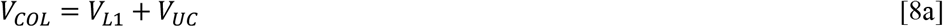

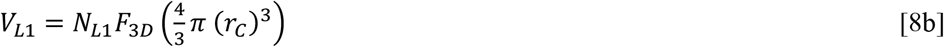

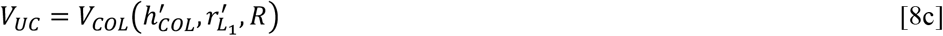

**Figure 3:**
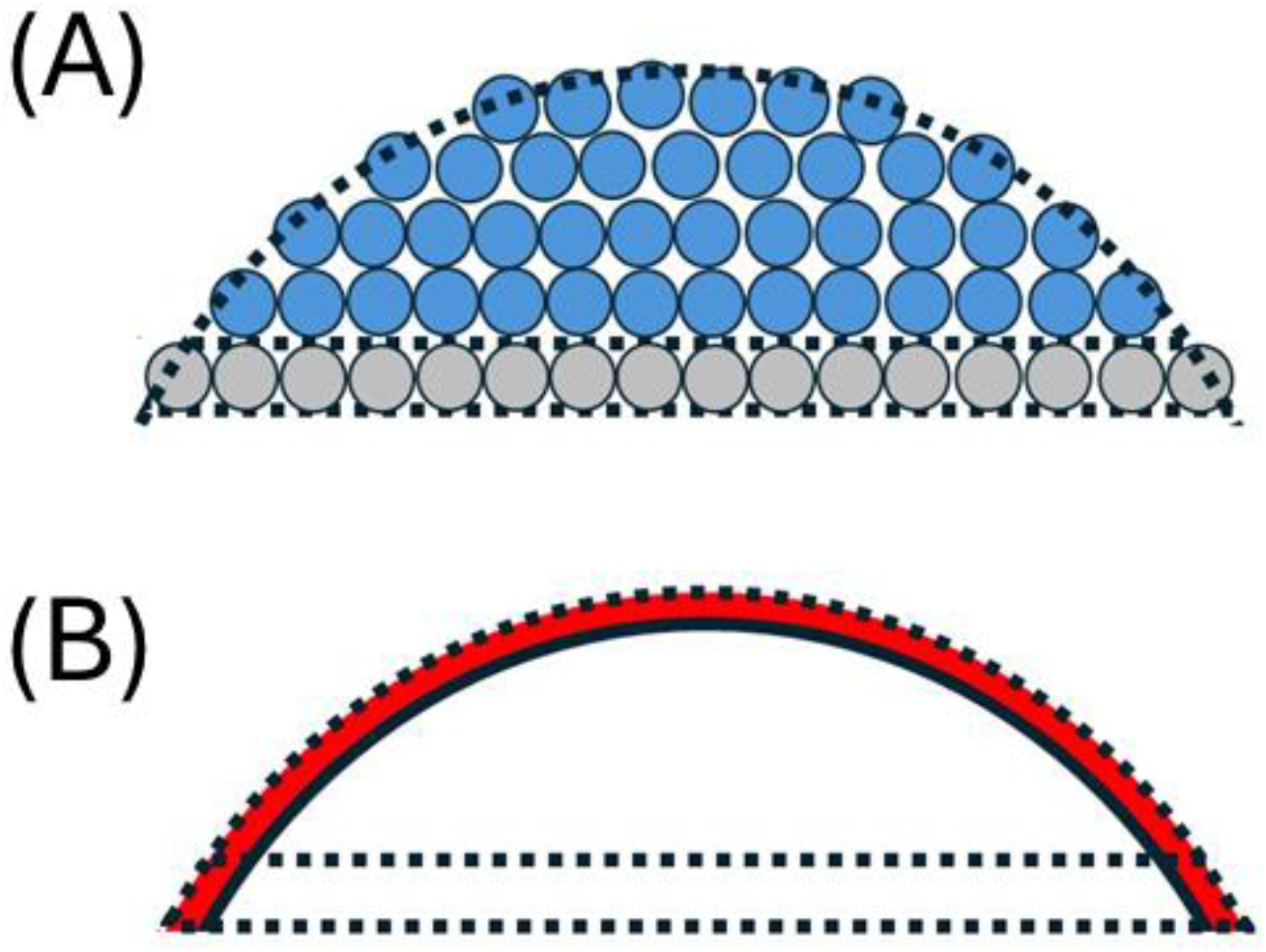
Sub-volumes of the spherical cap **(A) Total spherical cap volume, V**_**COL**_, **consists of the basal layer volume, V**_**L1**_, **and an upper cap volume, V**_**CAP**_ : Equations defining the relations between V_COL_, V_L1_ and V_CAP_ are given by Eqn. 7 and 8. **(B) Schematic description of internal and perimeter regions of two spherical cap sub-volumes:** Each sub-volume, V_L1_ and V_CAP_, consists of internal (white) and perimeter (red) regions with the number of cells in each region given by Eqn. 5c and d (basal layer) and Eqn. 9a and d (upper cap).

This geometrical conception of the cell colony, as a spherical cap partitioned into two sub-volumes, can be used to assign the number of perimeter and internal cells within the colony. Within the upper spherical cap volume the number of internal cells can be calculated through application of the functional form of Eqn. 7 with a reduced colony height, *h*_*COL*_*’’* (where 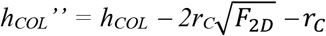), a reduced radius of the greater sphere *R’’* (where *R’’ = R − r*_*C*_) and a reduced radius of the circle of intersection, 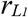 *’’* (where 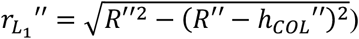 (**Eqn. 9a**) with the number of perimeter cells given by **Eqn. 9b**.

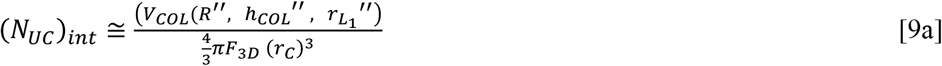

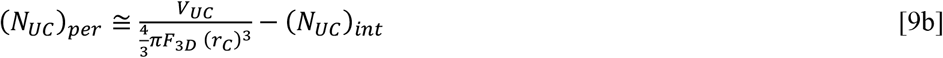

Calculation of the number of internal and perimeter cells within the basal cell layer is somewhat dependent on the biological effect of the surface on the growth of the cells directly in contact with it. Here we are principally concerned with the situation where the plate surface acts as a contact inhibitor of cell growth in the vertical direction (**Fig. 3b**)^4^. As the number of perimeter and internal cells contributing to lateral growth is accounted for within Eqn. 5e we need explicitly treat here only the number of basal layer perimeter and internal cells contributing to vertical growth which we designate as 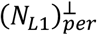 and 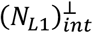. As this designation will be necessarily dependent upon the degree of coverage of layer one by layer two we utilize a transition function written in terms of the parameter *η* (where *η = 1−N*_*L1*_*/N*_*TOT*_) (**Eqn. 10 a-c**).

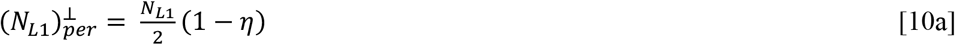

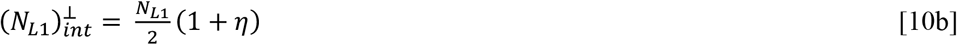

Evaluation of the number of perimeter and internal cells contributing to growth in both the basal layer and the upper cap allows for the calculation of the number of cells formed at each timepoint via appropriate numerical solution of **Eqn. 11**.

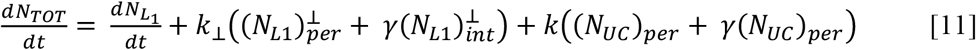

Joint numerical solution of the pair of inter-related ordinary differential equations defined by Eqn. 11 and Eqn. 5e respectively yield *N*_*TOT*_ and 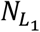 as a function of time along with the absolute dimensions of the growing colony.

### Simulation and characterization of growing colony

To assist with the interpretation of the cell colony geometry we examine the time evolution of parameters reflecting radius of the colony surface layer (*r*_*L1*_), the colony surface height (*h*_*COL*_), the radius of curvature of the colony (*R*), a composite shape term, *Ω*, (defined by Eqn. 12),^5^ and the incident contact angle between the surface and the colony edge, θ (defined by Eqn. 13).

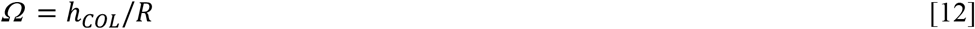

The contact angle θ is evaluated directly from the simulations according to Eqn. 13a and 13b which represent the respective cases when the spherical colony caps are smaller or larger than the equivalent hemisphere defined by the value of R.

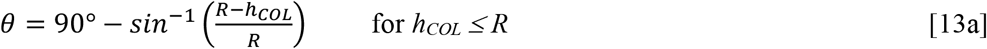

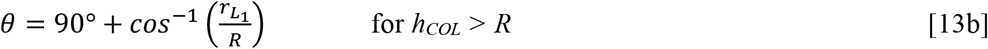

In all simulations we consider a spherical cell of diameter 2 μm with a doubling time of 90 minutes (translating into a lumped first order rate constant of k = 0.077 min^−1^) – values which are characteristic of bacteria and simple eukaryotes such as yeast **[Gray and Kirwan, 1974; Hartwell and Unger, 1977; Hall, 2023; Gaizer et al. 2024]**. For the completely isotropic case growth parameters were set as *k*_∥_/k = 2/3 and *k*_⊥_/k= 1/3 whilst for the horizontally and vertically preferred anisotropic growth cases the following parameters were respectively used; horizontally favored growth {*k*_∥_/k = 5/6 and *k*_⊥_/k = 1/6}; vertically favored growth {*k*_∥_/k = 3/6 and *k*_⊥_/k = 3/6}. For this set of parameter regimes the degree of anisotropy, Γ, defined by Eqn. 14, transitions from Γ = 0.5 to Γ = 2.5 between the three cases^6^.

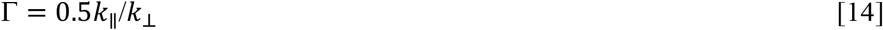

To produce a colony size sufficient to transition the microscopic/macroscopic measurement regimes we simulate for 3000 minutes which is slightly more than two days of culture – also values typical for cell culture / biofilm production experiments **[Nguyen et al. 2004; Huergo et al. 2012; Pokhrel et al. 2024]**. A constant packing fraction of F_2D_ = 1.2 (requiring F_3D_ = 1.32) reflective of imperfectly packed spheres was used throughout the simulations **[Scott and Kilgour, 1969; Kausch et al. 1971; Lubachevsky et al. 1991]**.

## Results

In describing the output of the model we first examine the rate of production of total cell number and the kinetic development of the geometric properties of the cell colony for the isotropic (**Fig. 4**) and anisotropic (**Fig. 5** and **Fig. 6**) cell division/growth cases, with each featuring five different extents of contact inhibition (over the range of γ= [0, 0.25, 0.5, 0.75, 1.0]). In comparing cell production over time (*cf*. Fig. 4A, 5A and 6A) we note that the effect of increasing the degree of contact inhibition (i.e. by increasing β and lowering γ) leads to a general decrease in the number of cells produced over time for both isotropic and anisotropic cases, with the very slight shape induced differences between the three cases disappearing in the limits of zero contact inhibition (γ = 1). The positive correlation seen between the reflective marker of contact inhibition, γ, and total cell number is also realized for both cases in the relation between γ and the markers of absolute colony size, denoting – the radius of the basal cell layer, r_L1_ (Fig. 4B, 5B and 6B), and the maximum height of the colony, h_COL_ (Fig. 4C, 5C and 6C). As revealed by the time evolution of shape parameter results reflecting the receding colony edge contact angle, θ (Fig. 4D, 5D and 6D) and the composite parameter, Ω (inset to Fig. 4D, 5D and 6D), changes in the degree of contact inhibition can have either a minor (in the isotropic case Fig. 4D or vertically favored anisotropic growth case Fig. 6D) or an extreme (for horizontally favored anisotropic case Fig. 5D) effect on the colony geometry. For the isotropic and horizontally favored anisotropic growth cases, greater extents of contact inhibition (increased β, decreased γ) eventually lead to more spherical bud-like colonies, counteracting the normal tendency to produce either near hemispherical growth (in the isotropic case Fig. 4D) or flattened colonies (in the horizontally favored anisotropic case Fig. 5D). However, this general trend is reversed for the vertically favored anisotropic case, with more perfect spherical buds being produced by lesser extents of contact inhibition (decreased β, increased γ). The general nature of the colony shape for the isotropic (Fig. 4E) and the respective horizontally (Fig. 5E) and vertically favored anisotropic cases (Fig. 6E) can be visualized from a progressive overlay of the outline of the central midsection (updated every 500 minutes) for the middle extent of contact inhibition (γ= 0.5 condition).

**Figure 4:**
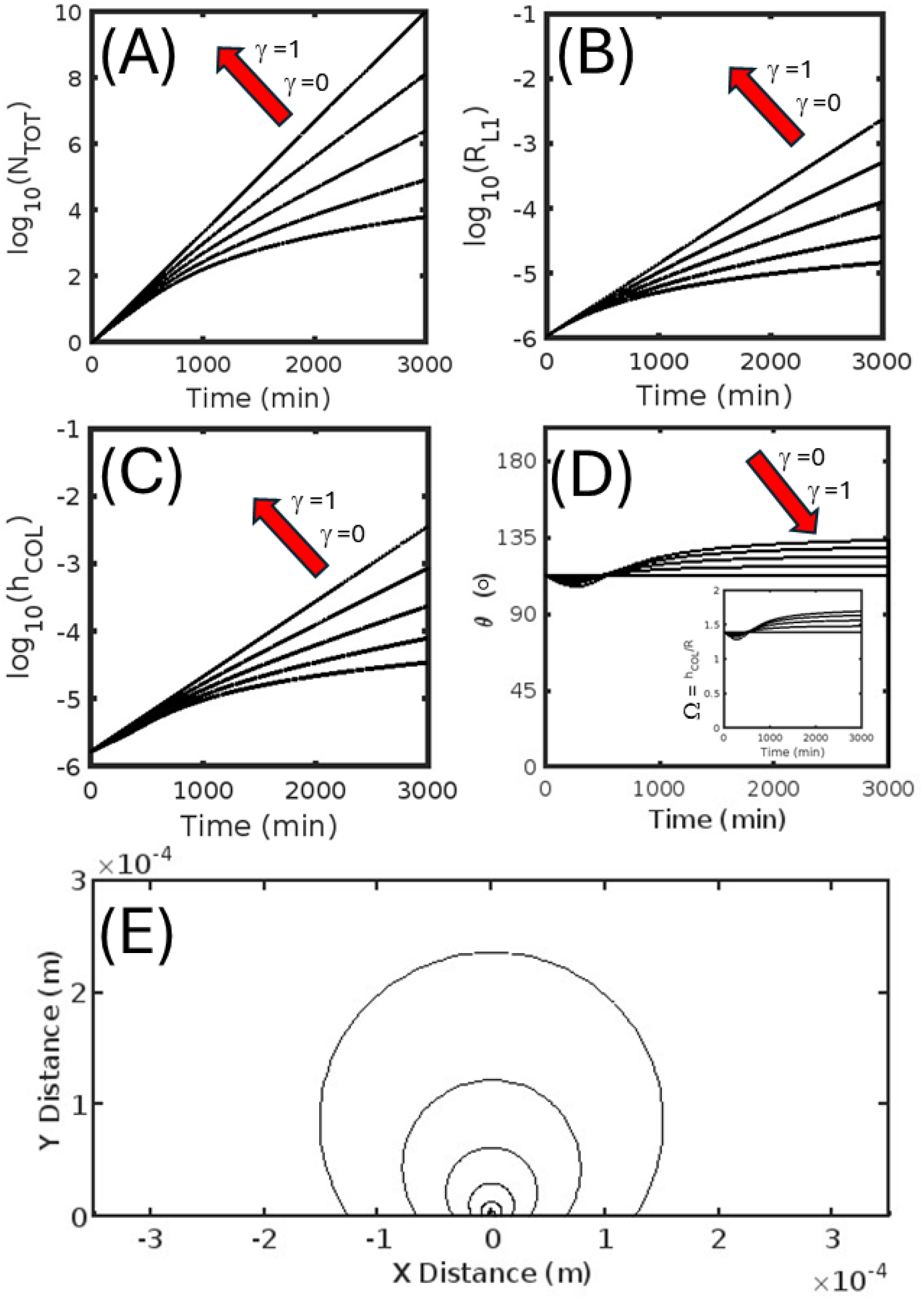
Isotropic colony growth (Γ = 1) occurring on a two-dimensional surface – Kinetic markers of total cell number and colony shape shown for five different extents of the contact inhibition parameter γ taken from the values [0, 0.25, 0.5, 0.75, 1.0]. **(A) Base ten logarithm of the total number of cells, N**_**TOT**_, **as a function of time. (B) Base ten logarithm of the radius of the colony basal layer, r**_**L1**_, **as a function of time. (C) Base ten logarithm of the colony height, h**_**COL**_, **as a function of time. (D) Receding contact angle, θ, of the colony as a function of time:** (INSET) Composite shape parameter, Ω, as a function of time. **(E) Colony longitudinal midsection profile:** Shown for six different times [500, 1000, 1500, 2000, 2500, 3000] mins.

**Figure 5:**
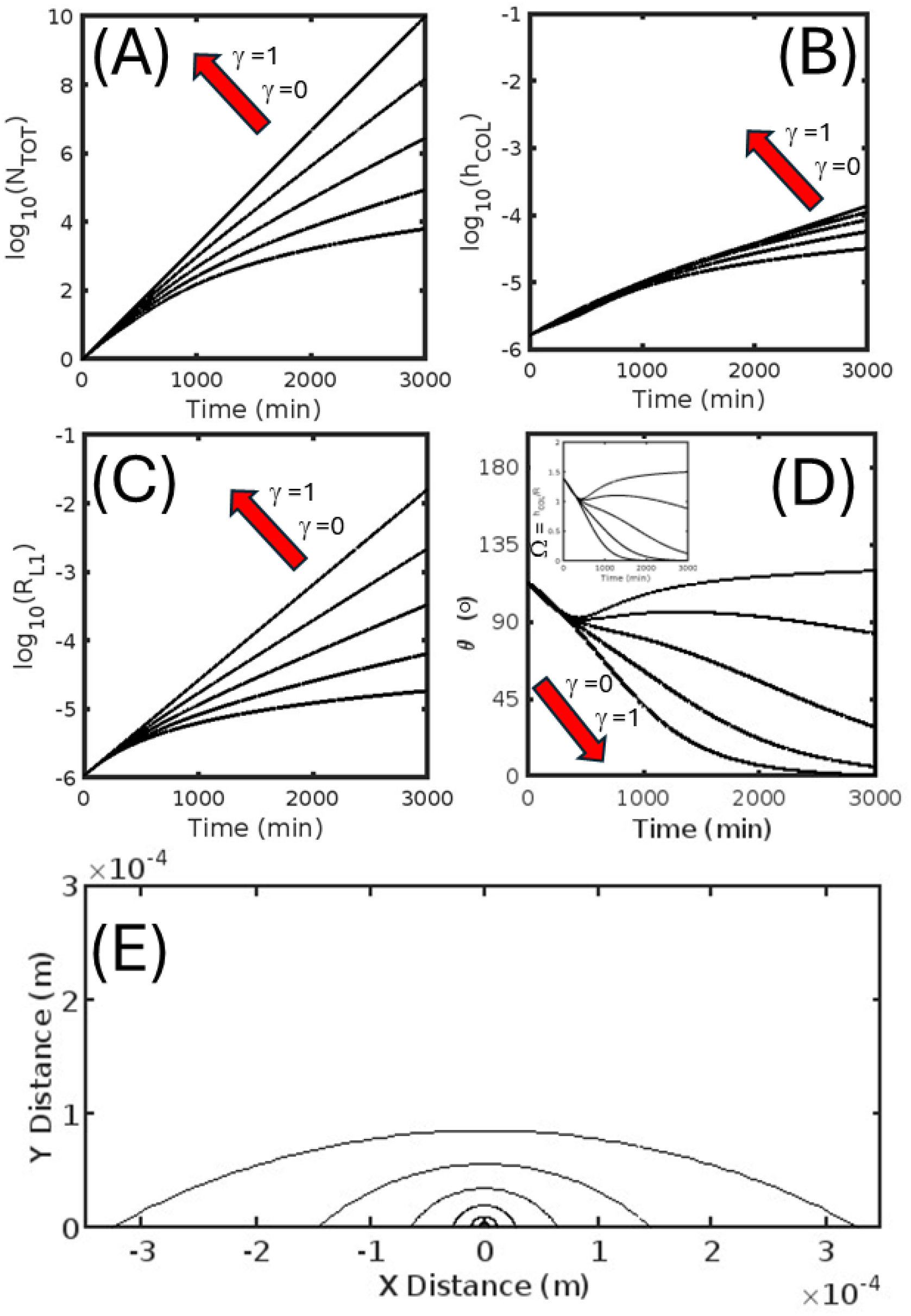
Anisotropic (laterally favored) colony growth (Γ = 2.5) occurring on a two-dimensional surface – Kinetic markers of total cell number and colony shape shown for five different extents of the contact inhibition parameter γ taken from the values [0, 0.25, 0.5, 0.75, 1.0]. **(A) Base ten logarithm of the total number of cells, N**_**TOT**_, **as a function of time. (B) Base ten logarithm of the radius of the colony basal layer, r**_**L1**_, **as a function of time. (C) Base ten logarithm of the colony height, h**_**COL**_, **as a function of time. (D) Receding contact angle, θ, of the colony as a function of time:** (INSET) Composite shape parameter, Ω, as a function of time. **(E) Colony longitudinal midsection profile:** Shown for six different times [500, 1000, 1500, 2000, 2500, 3000] mins.

**Figure 6:**
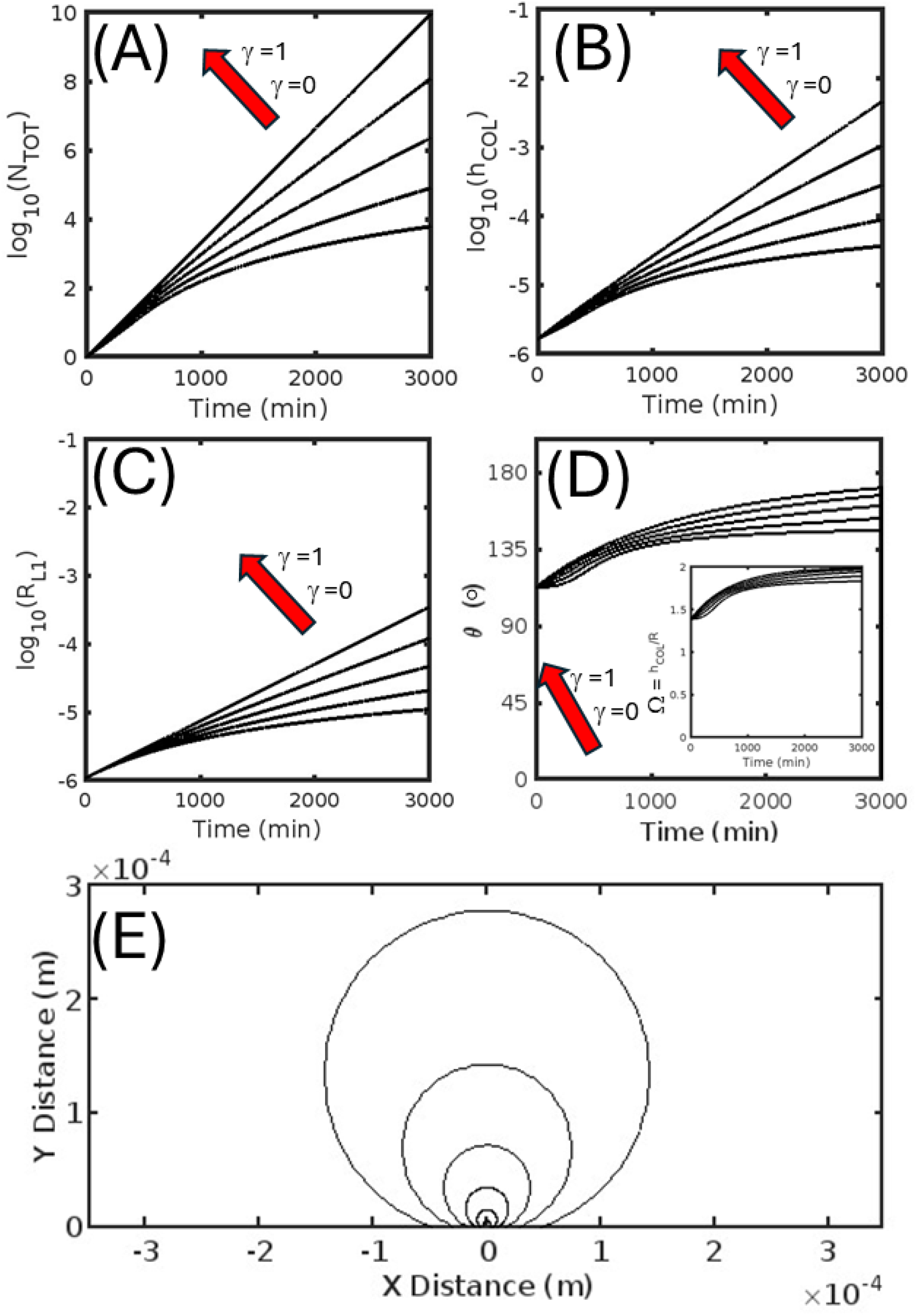
Isotropic colony growth (Γ = 0.5) occurring on a two-dimensional surface – Kinetic markers of total cell number and colony shape shown for five different extents of the contact inhibition parameter γ taken from the values [0, 0.25, 0.5, 0.75, 1.0]. **(A) Base ten logarithm of the total number of cells, N**_**TOT**_, **as a function of time. (B) Base ten logarithm of the radius of the colony basal layer, r**_**L1**_, **as a function of time. (C) Base ten logarithm of the colony height, h**_**COL**_, **as a function of time. (D) Receding contact angle, θ, of the colony as a longitudinal midsection profile:** Shown for six different times [500, 1000, 1500, 2000, 2500, 3000] mins.

## Discussion

In this paper, a macroscopic model of variable contact inhibited cell growth on a hard surface has been developed. To establish the potential limits of applicability of our model and to place the current work in both biological and physical context, we first provide a description of the basics of collective cell growth and the biological causes and consequences of contact inhibition. We then discuss the strengths and limitations of our physical modelling approach against those developed by others. We conclude with a discussion of the possible ramifications of the findings presented in this paper for the various scientific fields which involve the study of proliferative cell growth. Although the model developed in the present paper is most applicable to the description of eukaryotic cells cultured from multicellular organisms in the following discussion we will take a broad view of cell colony formation, extending it to both diverse types of cell growth and different causes of cell growth arrest, areas that are often treated separately but which feature sufficient commonality to generate insight when discussed collectively **[e.g cf. Shapiro, 1998 with Kamath and Bungay, 1988; Nguyen et al. 2004; Galle et al. 2005; Huergo et al. 2012; Hartmann et al. 2019]**.

### Mechanical aspects of colony growth

For most scientists, their first deliberative encounter with cell growth is via the culture experiment i.e. the controlled growth and division of cells under laboratory conditions **[Wistreich, 2002; Bonifacino et al. 2004; Jedrzejczak-Silicka, 2017]**. Although cell culture can be performed with cells from any of the three domains (archaea, bacteria or eukaryotes), irrespective of the type of cell used, all solid cell culture experiments have the same basic three requirements, the cell to be cultured, a growth surface or scaffold^7^, and a growth medium **[Wistreich, 2002; Bonifacino et al. 2004; Jedrzejczak-Silicka, 2017] (Fig. 7)**. Within this required framework there are three basic variants of what is known as a two-dimensional cell culture process **[Adams, 1990; Bonifacino et al. 2004]** which may be summarized as (i) cells plated on a hydrogel – typically used to culture bacteria and simple unicellular eukaryotes (such as yeast) in which cells are grown on top of a dense hydrogel loaded with liquid growth medium **[e.g. Nguyen et al. 2004; Pokhrel et al. 2024]** (Fig. 7A), (ii) adherent (or partially adherent) cells in a liquid/solid culture – a standard technique used to grow specialist differentiated eukaryotic cells isolated (or derived) from a multicellular organism, in which cells initially in solution, attach themselves to a plate surface by formation of protein mediated biochemical linkages known as cell adhesion molecules (CAMS) **[Lu et al. 1995; Nardone et al. 2011; Ahmad-Khalili and Ahmad, 2015]** (Fig. 7B), and (iii) cell growth concomitant with biofilm formation – a process in which cells (often subject to some form of nutrient limitation) collectively create a viscous film through the secretion of extracellular polymeric substances (EPS), that effectively enclose the growing colony promoting retainment of liquid and nutrients **[Su et al. 2012; Hartmann et al. 2019; Maier, 2021; Sauer et al. 2022] (Fig. 7C)**. With regard to the present paper, an important qualifying point to note is that the respective colony entry and exit points of nutrients and metabolic waste will be different for each of these three different types of two-dimensional cell culture experiment.

**Figure 7:**
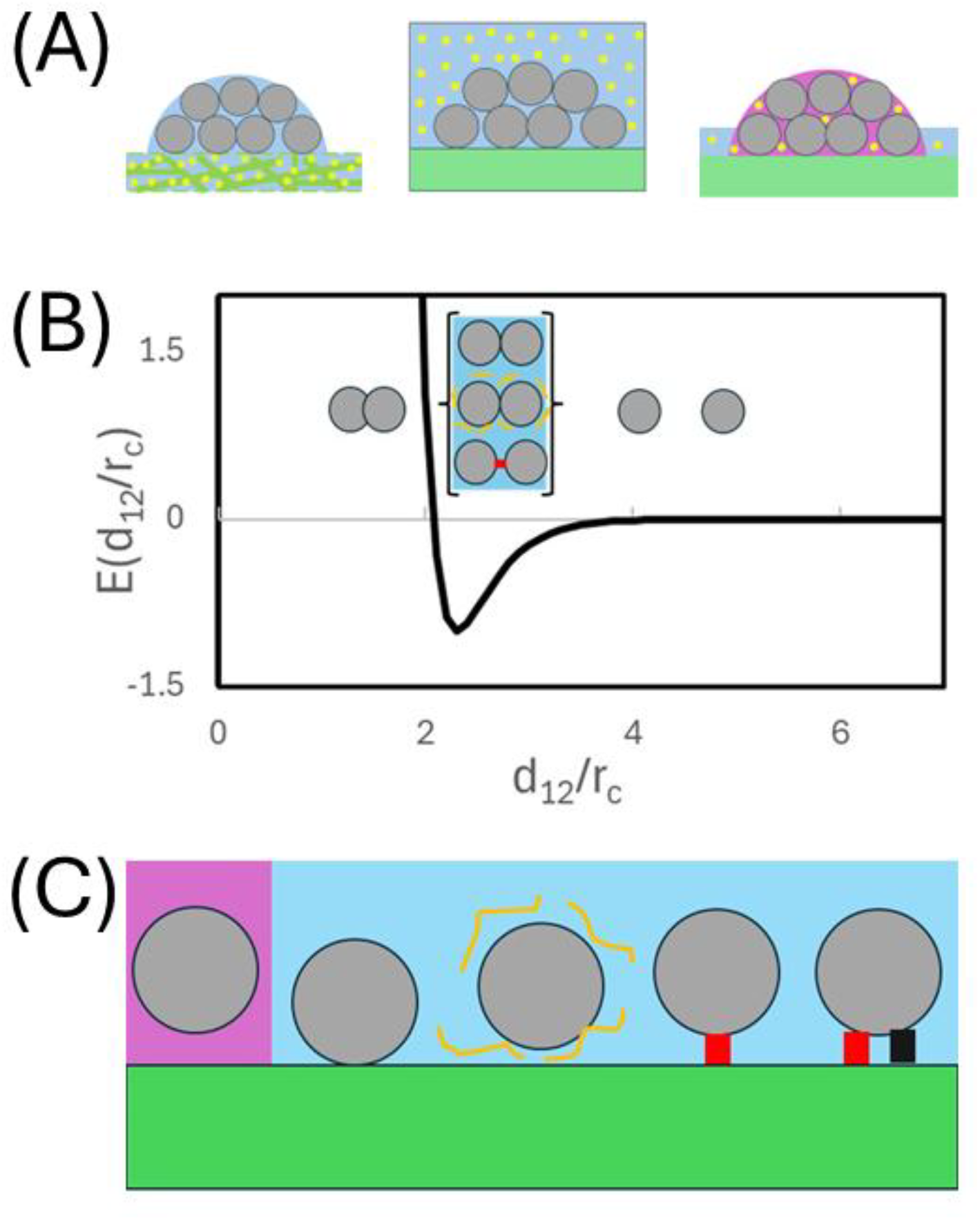
Viewing cell colony growth from a mechanical perspective. **(A) Three general formats of two-dimensional cell culture:** From left to right; Cells grow at an air/hydrogel interface in which the hydrogel is embedded with liquid and cell nutrients/food; Cells grow at a liquid/solid interface with nutrients/food contained within the liquid phase; Cells grow within a biofilm (hydrated viscous polymer) secreted by cells in response to a signal with the biofilm acting to store/concentrate liquid, nutrients and food. (Cells shown as grey circles, nutrients shown as yellow circles and metabolites shown as black diamonds). **(B) Schematic showing cell-to-cell attractive forces:** Intercellular attraction is indicated as a drop in reduced energy of the system, E*, as function of cell-to-cell distance normalized with respect to spherical cell radius (d_12_/r_C_). Three attractive/repulsive regimes are shown; steric repulsion below the intercellular contact distance; weak to strong attraction at or just beyond the intercellular contact distance promoted by various direct (covalent coupling, hydrogen bonding, ionic interactions) and indirect (e.g. hydrophobicity, polymer depletion, polymer/protein mediated interaction) forces; Non-attractive regime where the cells are separated by an appreciable distance. **(C) Schematic showing cell-to-surface attractive forces:** Similar to the case described above for cell-to-cell interactions, cell-surface interactions can be promoted via direct or indirect forces or by various types of specialist surface anchoring proteins known as cell adhesion molecules.

Although the experimental process differs among each of these three types of solid phase two-dimensional cell culture formats, in each case the cells within the growing colony are effectively bound to each other, and to the surface, by attractive forces of chemical / biochemical origin (Fig. 7D and 7E) with gravity playing little role in dictating the behavior of an isolated cell^8^. As discussed by Maier **[Maier, 2021]** (and summarized within Fig. 7D) cells will generally repel each other at distances less than that of closest approach due to steric exclusion **[Maier, 2021]**. Conversely, cells will attract each other at, or slightly greater than, the contact distance, via a range of possible forces/pseudo-forces mediated by localized hydrophobicity/electrostatic interactions, colloidal double layer effects, polymer depletion and specific protein-mediated linkages (such as are manifested in occluding junctions, anchoring junctions and communicating junctions)^9^ **[Maier, 2021]**. As summarized in the schematic (Fig. 7E) cell attachment to surfaces or polymer matrices can likewise be effected through nonspecific interactions with the surface **[Maier, 2021]**, nonspecific interactions with the components of an extracellular polymer substances (EPS) e.g. polysaccharides, proteins, lipids, and extracellular DNA **[Sauer et al. 2022]** or specific biochemical linkages such as those grouped within the cell adhesion molecules (CAMS) e.g. integrins, cadherins, immunoglobulins, selectins, and specialist proteoglycan receptors **[Ahmad Khalili and Ahmad, 2015]**.

Due to the cohesive nature of cells to both each other, and to the surface, any cell division and growth occurring internally within the colony will call it to swell and change its overall shape. An early model for predicting cell colony geometry used surface tension arguments suitable for describing the shape of liquid droplets adsorbed to an interfacial boundary **[Reuter and Taylor, 2006; Nguyen et al. 2004]**. Although providing useful physical insight such approaches suffered from the demonstrable non-liquid nature of cell colony which is more closely approximated by a semi-elastic irregular gel **[Constantinides et al. 2008]**. Within such an elastic material framework, any structural deformation of the colony will be defined by the point of origin of the expansion occurring within it, the vectorial nature of the force exerted at the point of expansion, and the material elasticity tensor describing the response of the colony to the exerted force **[Byrne and Drasdo, 2009; Chaplain et al. 2020]**. The material elasticity tensor could, in principle, be determined from analysis of measurements of resistance to direction specific pushing or pulling forces applied to the colony using magnetic or optical tweezers **[Catala-Castro et al. 2022]** or atomic force microscopy experiments **[Kosheleva et al. 2023]**. In the absence of an external force, cell growth within a material defined by uniform and isotropic elasticity tensor, will result in a spherical growth cap colony geometry for the cases of (i) isotropic cell growth, and (ii) anisotropic cell growth with subsequent capability for reorientation/relocation of the dividing cells as a means of minimizing the potential energy of the stretched colony **[Nguyen et al. 2004; Byrne and Drasdo, 2009; Chaplain et al. 2020]**. Such an understanding sets a strong requirement for the existence of either the existence of an imposing external force (e.g. gravity or an applied shear force) or a non-isotropic elasticity tensor for the colony (i.e. different degrees of stretch for the same magnitude of force applied in different directions) to produce a colony shape different from a spherical cap. Within this type of mechanical framework (i.e. that of a semi-elastic gel) the colony shape will be affected by the presence of external forces (such as liquid shear and gravity) with the inclusion of these effects becoming more important as the colony becomes larger **[Warren et al. 2019; Pokhrel et al. 2024]**. In the absence of detailed mechanical measurements of the direction-dependent deformability of cell colonies it is tempting to speculate that the reasons why colonies are frequently observed to adopt a spherical cap structure in the early stages of their growth is due to the isotropic elastic properties of the colony ‘gel’. It is also interesting to recognize that a common means for changing the local elasticity within gels is via their partial dehydration at their surface exposed edges **[Kerch, 2018]**.

### Biochemical aspects of contact inhibition

In order to grow and reproduce cells require an energy source and, like all machines, they generate both waste and chemical byproducts as a result of the energy usage/conversion process (**Fig. 8A**) **[Adams, 1990]**. Among both simple and complex cell types, a range of substrate induction and product inhibition feedback mechanisms exist to regulate both the occurrence and pace of cell division and growth **[Ward and Thompson, 2012; Zhu and Thompson, 2019; Basu et al. 2022]**. As a result, even when existing in a solitary planktonic state, cell growth and division can be differentially regulated by nutrient surplus or limitation and the presence of excess waste metabolites **[Monod, 1949; Hartwell and Unger, 1977]**. In addition to their activation/deactivation by metabolites, nearly all cells are capable of both sending and receiving specialist biochemical signals to/from other nearby cells to help induce and regulate advantageous collective growth behaviors (e.g. dormancy in response to material deficit, differentiation in response to colony / tissue development requirements) **[Water and Bassler, 2005; Zhu and Thompson, 2019]**. Such intercellular signaling can result from the direct intake of a chemical messenger, or a secondary process involving the interaction of a chemical messenger with a membrane receptor which in turn stimulates a transduction event capable of activating an internal ‘second messenger’ cascade **[Ward and Thompson, 2012]**. The signaling process can be even more convoluted, involving a form of auto-regulation occurring as the downstream result of an accumulation of prior changes in the cell, with one pertinent example being the cessation of the cell division cycle as a result of the formation of numerous intercellular and/or cell-surface linkages of the type described in the previous section **[Ribatti, 2017; Mendonsa et al. 2018; Alberts et al. 2022; Basu et al. 2022]**. In terms of the current work, whatever the mechanistic cause (whether it be nutrients, signaling molecules, or downstream regulation as a result of cell/tissue maturation – Fig, 8A) the eventual biochemical response results in the switching on or off (or partial tuning) of the cell division and growth process. Although somewhat cursory, this short description highlights the various major causative pathways of contact inhibition – a more detailed description of the specific cell cycle proteins (cyclins) is given in the appendix of the first paper in this series **[Hall, 2024]** and in the work by Basu et al. [**Basu et al. 2022]**.

In considering how these just discussed effectors of growth regulation (i.e. nutrient concentrations, metabolite/signaling molecule concentrations and cell-to-cell/ cell-surface connections) might be spatially manifest within a colony, one might venture the following general argument based on relative cell positions (**Fig. 8B**). For cells exhibiting an equal rate of cell growth and reproduction, those within the regions toward the interior of colony would over time likely experience lower concentrations of nutrients (due to their preferential accumulation by cells closer to the colony surface) and higher concentrations of excreted waste and secreted chemical signals (due to the more tortuous path required for their exit from the colony). Assuming a regular packing geometry, we might also suppose that cells existing at the colony edge region would possess a lower number of cell-to-cell contacts due to their having a lower number of cells in their direct proximity^10^. A simple example of how these qualitative arguments may be spatially manifest is provided (**Fig. 8C**). However, whilst such explanations share a basic reasonableness, translating qualitative speculations into quantitative predictions requires a more complex model of cell growth and division coupled with an often unreconciled level of system definition i.e. spatiotemporal description of nutrient, metabolite, signaling molecule concentrations within the colony, spatiotemporal description of cell structure, cell-to-cell connectedness and local elasticity, and spatiotemporal description of metabolic activity and growth and division rates. Although this point will be addressed more fully in the next section on mathematical models, it is worth noting that any cell colony modelling approaches that feature nutrient diffusion into a colony do so with limited knowledge of the relevant transfer and uptake rates, and are also often formulated without consideration of other potential regulatory mechanisms of cell growth/division induced by metabolic waste egress from the colony, cell signaling or cell-to-cell connections **[Ward and Thompson, 2012; Ahmad Khalili and Ahmad, 2015; Mendonsa et al. 2018]**. As a general point, it is entirely possible that the potential causes of contact inhibition will be ranked differently in importance for different cell types, modes of colony growth and concentrations of nutrients with these rankings also possibly changing over the time course of single experiment due to changing colony size and dimensions, and build up/depletion of waste and nutrients **[Ward and Thompson, 2012; Lavrentovich et al. 2013; Mendonsa et al. 2018]**.

**Figure 8:**
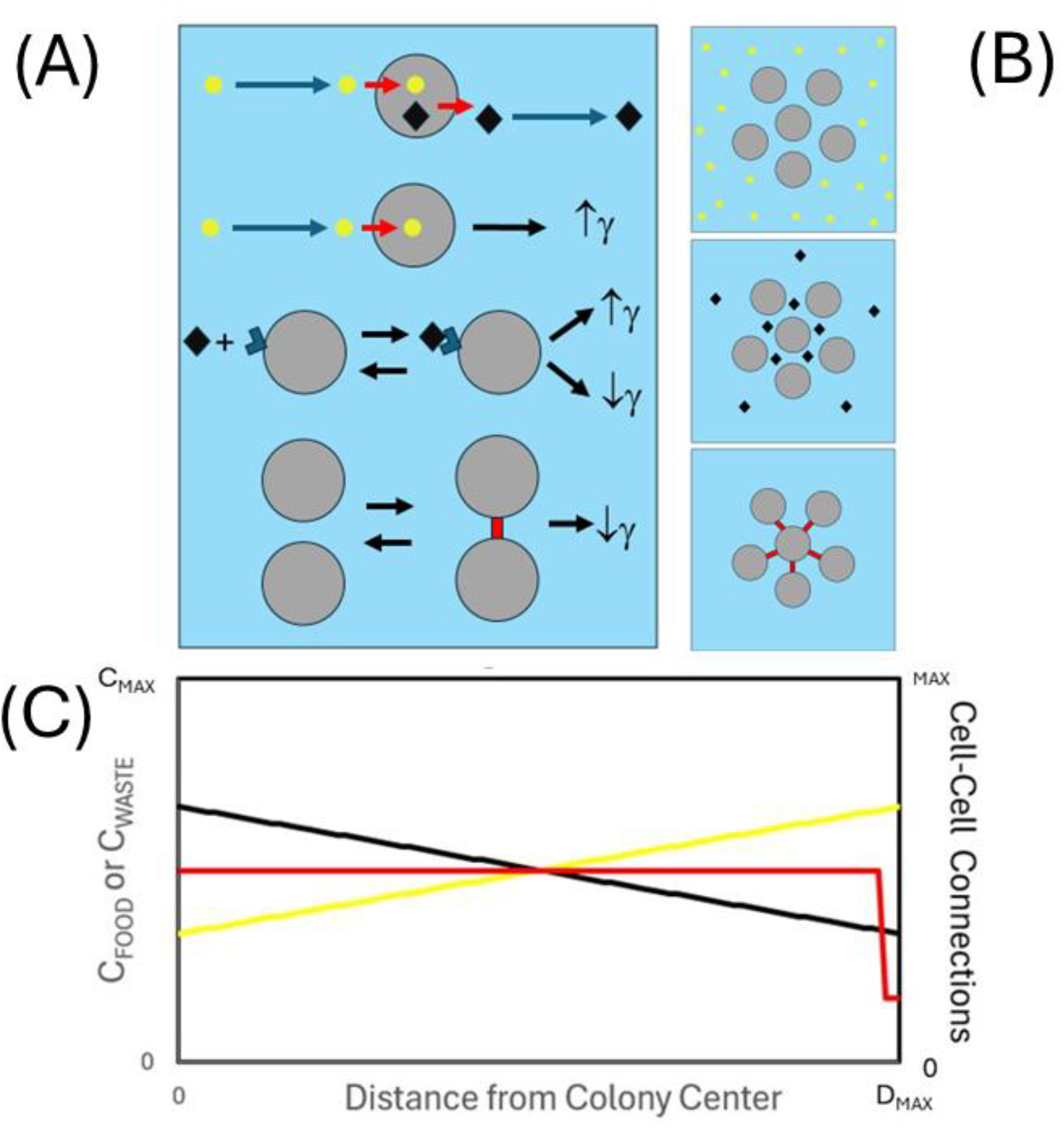
Viewing colony growth from a biochemical perspective. **(A) Schematic showing promoters and inhibitors of cell growth and division:** Living cells require nutrients and food sources and, as a result of metabolism, produce waste and other metabolites. In a general manner we might consider food sources as promoters of cell growth/division, metabolites/signaling molecules as either potential promoters or inhibitors of growth/division, and cell-to-cell contact as a potential inhibitor of cell growth/division. (Cells shown as grey circles, nutrients shown as yellow circles and metabolites/signaling molecules shown as black diamonds, cell-to-cell linker molecules shown in red). **(B) Schematic showing spatial considerations of nutrient/waste molecules ingress and egress from the colony along with potential for formation of cell-to-cell contacts. (C) Graphical example of a potential spatial profile of nutrient (yellow), metabolite (black) and number of cell-to-cell connections (red) as a function of distance from the colony center for a colony immersed in a uniform liquid bath.** Note that the situation will be quite different when the cell colony will grow at a surface for the three different cases shown in Fig. 7A.

### Limitations and strengths of the model in comparison with published literature

In this section a brief recount (sans equations) of what was done in the present work is outlaid, before contrasting the strengths and weaknesses of the model against approaches developed by others.

#### Current work

The current paper extends a previously developed macroscopic model of contact inhibition applicable to completely symmetrical cell colony growth **[Hall, 2024]** in order to treat the case of colony growth occurring on a flat surface. The current model is capable of accounting for variable anisotropy in the favored direction of cell growth/division within the basal layer, and a variable degree of contact inhibited growth/division based on whether cells exist at the colony edge or internally. Similar to prior models developed for completely symmetrical cases of spheroids and circular monolayers **[Radszuweit et al. 2009; Montel et al. 2013; Hall, 2024]**, the model developed in this work is internally consistent and continuous over all possible ranges (and therefore capable of accepting a single cell on the plate as the initial condition). The output of the model yields both the number of cells, and the colony dimensions, as a function of time. The model is based on three assumptions which we interrogate here in more detail.

#### (i) Directionality of the cell division/growth

The lumped cell division/growth rate constant was considered capable of being spatially resolved into its vector components (Eqns. 3 and 4). This assumption allowed for specification of the lateral versus vertical growth rate within the surface layer and as a result provided a means for partial specification of anisotropic growth tendencies. Although represented as simple spheres, bacterial cells are frequently asymmetric (e.g. E.coli bacteria are spherocylindrical **[Rudge et al. 2012; Beroz et al. 2018]**, and eukaryotic cells often display an irregular flattened shape **[Hall, 2023; Pincus and Theriot, 2007; Holmes and Edelstein-Keshet, 2012;]**). Cells may also make physical or biochemical changes in response to an asymmetric environment (e.g. epithelial cells may exhibit a polarity upon forming attachments to a surface **[McGarry and Prendergast, 2004]**, cells may change shape when existing within a nutrient or signaling gradient **[Dasbiswas et al. 2018]**). In the present model the surface is considered as the principal propagator of directional anisotropy effects. Growth/division anisotropy effects are considered to decay to zero for cells at some distance from the surface due to either newly formed cells possessing a lower sense of directionality (e.g. as associated with the buckling transition seen in certain types of asymmetric cell types **[Su et al. 2012; Beroz et al. 2018]**), or bulk material stretch-induced forces causing rearrangement/reorientation of cells following their division and growth **[Nguyen et al. 2004]**.

#### (ii) Base cell layer grows as a circle due to in-plane growth

A large number of experimental and simulation studies have shown that under standard growth conditions cells generally form circular colonies irrespective of whether the colonies are simple monolayers or extend into the vertical dimension **[Pirt, 1967; Kamath and Bungay, 1988; Galle et al. 2005; Huergo et al. 2012; Pokhrel et al. 2024]**. Within the interior of a colony featuring a significant vertical dimension, cells may come to exist at a position due to division processes occurring above or below or to either side. The assumption made in the current paper, that cell growth within the basal layer occurs principally from in plane growth, obviated the need to estimate the contribution to basal layer growth from cell division processes occurring above it. This assumption is likely relatively strong, indeed Newton’s third law implies that any downward force exerted by upper cells will lead them to be forced away from the surface. However, due to the existence of different types of cell-surface vs. cell-cell linkages, the possibility exists for both, different levels of adhesion between cells and between cells and surface, and different extents of induced contact inhibition i.e. (γ)_basal layer_ vs. (γ)_colony_ [see **Galle et al. 2005**]. Although not explored in the current paper, such effects would effectively constitute an equivalent form of growth anisotropy and would be expected to affect colony shape in a manner similar to that demonstrated in Figs. 4 to 6.

#### (iii) Colony shape modelled as a spherical cap

A spherical cap model was selected to approximate the growing cell colony shape on the twin basis that (i) significant empirical evidence suggests that many different types of cultured cells adopt this geometry for significant portions of their growth as a collective **[Palumbo et al. 1971; Wimpenny, 1979; Kamath and Bungay, 1988; Nguyen et al. 2004; Pipe and Grimson, 2008; Beroz et al. 2018; Warren et al. 2019; Balmages et al. 2023; Pokhrel et al. 2024]**, (ii) mechanical arguments imply that the assumption of a spherical cap geometry requires a uniform colony elasticity and, vice versa, the assumption of uniform colony elasticity specifies a spherical cap geometry **[Nguyen et al. 2004; Reuter and Taylor, 2006; Beroz et al. 2018]**. Consideration of these two points provides a means for understanding, rather than justifying, the predicted differences in colony shape for the two extreme anisotropic cases (**Fig. 5E** vs. **Fig. 6E**). An important requirement for satisfying the assumption of constant material properties is the existence of an effectively isotropic rate of cell growth and division throughout the internal region of the upper cap part of the colony. In addition to the mode of contact inhibition invoked within the current model, an alternative means for regulating cell/growth rate in a position dependent manner, is via the development of spatiotemporal concentration gradients of nutrients, waste and signaling molecules, in the regions within and around the cell colony **[Pirt, 1967; Kamath and Bungay, 1988; Vulin et al., 2014; Lavrentovich et al. 2013; Warren et al. 2019; Pokhrel et al. 2024]**. In some of the earliest studies of their kind, diffusion of both glucose and oxygen into a yeast colony growing on an agar plate were measured and shown, under conditions of limited supply, to become depleted within central regions of the colony and at adjacent regions of the agar plate, creating so called solute ‘depletion zones’ for cases of very large colony sizes (greater than mm internal diameter size) **[Pirt, 1967]**. Indeed, Kamath and Bungay **[Kamath and Bungay, 1988]** have used such arguments to rationalize why very large colonies transition from a spherical cap to a spherical cap napkin-like (SCNL) structure-a phenomena recently re-discovered through geometrical scaling analysis of seven types of bacterial growth **[Pokhrel et al. 2024]**. Others have noted that the deviation from a spherical cap structure at large colony size may result directly from changes in the colony gel properties **[Nguyen et al. 2004]**. Clever experiments based on coating the culture plate agar surface with a patterned mesh featuring regions of different nutrient permeability were used to further explore factors affecting yeast colony shape **[Vulin et al. 2014]**. When yeast cell growth was restricted to a porous circular area of 1.5 mm diameter, the growing yeast colony transition from hemispherical to columnar geometry in which growth of cells in the upper region of the column was severely retarded in a manner indicating nutrient transport limitation. It is interesting to speculate both on the limiting radial size dependence of this transition and also the yeast strain dependence (particularly with regards to strains exhibiting over or under expression of the yeast adhesion protein FLO1 e.g. see **[Nguyen et al. 2004]**). Similar experimental systems used to fabricate tumor spheroids or organoids from eukaryotic cell colonies **[de Carli et al. 2021]** indicate quite different modes of cell-to-cell adhesion.

#### Other modelling approaches

Due to the fact that colony formation by unicellular organisms represents the first basic step in the development of a multicellular organism **[Bassler and Losick, 2006; Ros-Rocher, 2021]** the simulation of the kinetics of colony and tissue development has received significant prior attention. Taking a historical perspective, here we review relevant previous research, dividing it into three areas, (i) continuum models, (ii) agent-based models, and (iii) geometrical scaling-based models. Each introduced work will be used to compare and critique the good and bad aspects of the current work.

#### (i) Continuum models

This type of model typically describes the number of cells, the colony volume and the overall colony shape using a set of ordinary or partial differential equations, concentrating on bulk properties with either no information given on the behavior of individual cells **[Lowengrub et al. 2009; Chaplain et al. 2020]**. A continuous model of contact inhibited spheroidal tumor growth based on a single ordinary differential equation was first developed by Mayneord **[Mayneord, 1932]** with improvements to this general approach subsequently made by others **[Radszuweit et al. 2009; Montel et al. 2012; Alessandri et al. 2013; Hall, 2024]**. For completely symmetrical structures such as circles (for monolayer growth) and spheres (three-dimensional growth) colony size scales directly with cell number making prediction of the absolute colony dimensions straightforward. An early attempt at modelling bacterial colony growth on a hard surface was made by Pirt who, by treating colony shape using a fixed hemisphere approximation, could also employ the direct scaling relations between colony size and cell number made for the completely symmetrical cases of spherical and circular colony growth [**Pirt, 1967**]. One important aspect of the work by Pirt was the mathematical rationalization of the frequently observed transition from an exponential to a linear growth rate of both cell colony radii and height amongst many different types of cultured cells [e.g. cf. **Pirt, 1967; Kamath and Bungay, 1988; Huergo et al. 2012**; **Bravo et al. 2023**] by considering a ‘growing’ zone of cells located at the colony periphery and a ‘non-growing’ zone of cells located at the colony’s internal regions with the dimensions of the growing zone shown to be sensitive to nutrient (glucose and oxygen) concentration. A more sophisticated continuous version of this two-state model has been shown to describe linear growth rate of biofilm height for many different kinds of bacteria **[Bravo et al. 2023]**. Such linear growth behavior in both radius and height is also predicted in the present work, showing up as significant curvature in the log_10_(r_L1_) and log_10_(h_COL_) vs time plots shown in Figs. 4B/C, 5B/C and 6B/C. Further experimental and theoretical work finessed this approach by approximating growing bacterial and yeast colony shapes using a spherical cap model, empirically defined by their experimentally recorded circular base radii and colony heights **[Palumbo et al. 1971; Kamath and Bungay, 1988]**^11^. More recently, an approach using a partial differential (position and time dependent) formulation of colony growth in which internal pressure gradients generated by cell division and growth are used to evaluate the colony shape directly from the simulation (subject to capability of adequately specifying the required parameters) **[Greenspan, 1976; Byrne and Drasdo, 2009; Chaplain et al. 2020]**. A more modest partial differential equation-based approach requiring assumption of a fixed geometry has been developed for the simulation of the diffusion limited growth of *E. coli* spheroids grown in soft agar **[Shao et al. 2017]**. This latter approach coupled with a spherical cap approximation of the colony geometry could provide a more detailed description of colony surface growth subject to nutrient limitation but, similar to the present work, in the absence of a constraining shape approximation, it has limited ability to predict colony shape de novo.

#### (ii) Agent-based/discrete models

These types of models attempt to specifically account for each cell within the colony, representing them explicitly (e.g. by use of an index or shape with assigned properties) and allowing them to interact with other cells through a defined set of specified rules^12^ **[van Liederkerke et al. 2015; Picioreanu et al. 1998; Montagud et al. 2021]**. The first agent-based model of cellular growth described two-dimensional monolayer formation **[Eden, 1960]** and since then similar, but progressively more sophisticated, approaches have been used to describe two-dimensional surface growth **[Kreft et al. 1998; Rudge et al. 2012; Aland et al. 2015; You et al. 2018; Schnyder et al. 2020]**, spheroid growth **[Radsuweizet et al. 2009; Montel et al. 2011; Waclaw et al. 2015]**, and three-dimensional colony formation at a hard surface **[Galle et al. 2005; Su et al. 2012; Beroz et al. 2018; Hartmann et al. 2019; Warren et al. 2019]** with this latter class of models tending to yield spherical cap-like structures (e.g. cf. [**Galle et al. 2005; Beroz et al. 2018; Hartmann et al. 2019]** vs. Warren et al. who approximated colony growth as a triangular cone **[Warren et al. 2019]**). Similar to results reported from the continuum models (described in the previous section) and the results obtained in the present paper, all of the cited agent-based models demonstrate an effective transition from exponential to linear colony radius and colony height growth rates over the time course of colony expansion with some studies showing interesting changes in shape during the colony growth time course (e.g. see Fig. 4 of **[Beroz et al. 2019]** and Fig. 1H of **[Warren et al. 2019]** and compare to Fig. 4D, 5D and 6D in the current work).

#### (iii) Geometrical scaling models

An alternative approach to quantitation is the empirical description of experimentally derived characteristic points, sometimes called ‘model free’ analysis. When the processes being studied follow some fundamental rule or convention data taken from different experiments can often be collapsed onto a common dependence by performing a suitable scaling arrangement **[De Gennes, 1979]**. With regards to the kinetic evolution of cell colony geometry a number of studies have found such scaling relationships to varying degrees **[Pirt, 1967; Kamath and Bungay, 1988; Pipe and Grimson, 2008; Bravo et al. 2023; Pokhrel et al. 2024]**. Perhaps the most basic model free finding is that cell colonies often adopt a spherical cap-like geometry during much of their early time course with cap section radius and colony height initially evolving with an exponential dependence on time which then later becomes a linear dependence [**Pirt, 1967; Pipe and Grimson, 2008; Bravo et al. 2023; Pokhrel et al. 2024**].

### Relevance of the current findings to fields involving proliferative cell growth

The model developed in the current paper has some potential relevance beyond its use in the direct quantitation of different types of cell colony growth. Of potential interest to those involved with cancer research is the dramatic change to tumor shape signaled by an alteration to the growth anisotropy characteristics of the tethered layer (Fig. 5D). Indeed, a sudden change in lateral growth preference of cells in a basal layer to isotropic/vertically favored growth would manifest as a planar to spherical transition in local tumor shape **[Lu et al. 1995; Nardone et al. 2011]** (Fig. 5D) – potentially providing a rather simple physical explanation of one of the early stages of metastasis i.e. the process by which smaller tumors can detach from a primary cancerous growth prior to their spread/circulation through the body. Also relevant to cancer biology is in the model’s capability to empirically assign an effective ‘respect of confluence parameter’ (γ) to cells grown in culture. The assignment of a value of γ, together with the standard proliferation rate, k, may provide a more complete quantitative indicator of the cell’s cancerous nature under a particular set of conditions. Conversely, this parameterization strategy may assist with direct assessment of colony structure in cell culture-based testing of (i) anti-cancer drugs – such as those intended to disrupt the tubulin/microtubule polymerization pathway **[Imamura et al. 2015; Hall, 2003]**^13^ or (ii) cell toxins – such as the lysogenic amyloid fibers **[Solomon, 1997; Hall and Edskes, 2004; Hall and Edskes, 2009; Sasahara et al. 2013]**.

## Conclusions

Although the power of mathematics often lies in its potential for abstraction and generalization, there is always the risk that any general model will be deemed unsuitable to address the complexities of a particular biological case. In this paper we have developed a continuum model of cell colony growth on a hard flat surface that unlike recent quantitative efforts based on scaling analysis of measured values of colony surface area and height **[Pokhrel et al. 2024]** allows for an internally consistent simulation approach to the evolution of the growing colony shape as a function of time based on just two characteristics, the cell growth rate matrix, **k**, and the cell confluence parameter, γ. The model uses a strict conceptualization of the contact inhibition phenomenon by defining regions of non-contact inhibited cells at the colony surface and partially contact inhibited cells within the colony’s internal region. Due to its minimal parameter requirement and its ability to describe shape as well as volume the approach also has potential for use in data reduction of various types of cell colony growth assays. Importantly, the model can accommodate transitions from monolayer to multilayer growth via alteration of a single parameter.

## Acknowledgements

I would like to thank Prof. W. K. Olson for helpful scientific discussions and for comments made on an early draft of this manuscript. Thanks to Mr. A. Richard and Ms. J. Ann for a careful reading of the manuscript and for providing suggestions for improvements in the overall clarity of expression. I gratefully acknowledge support from Rutgers, State University of New Jersey, provided in the form of a Visiting Scientist appointment held within the Department of Chemistry and Chemical Biology, Center for Quantitative Biology.

## Author Contributions

Damien Hall was responsible for conceiving, performing and writing the work.

## Appendix 1

Dependent upon the exact biological mechanism of the cell contact inhibition the potential exists for an alternative situation to the one considered in the main text in which the plate surface does not act as a contact inhibitor of cell growth in the vertical direction (**App. 1 Fig. 1a**). In this case the bottom facing component of the basal layer will always constitute a perimeter region with the calculation of 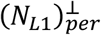 and 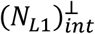. given by Eqn. App. 1-1 a and b.

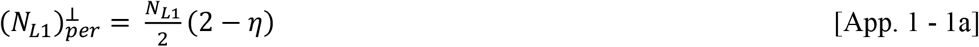

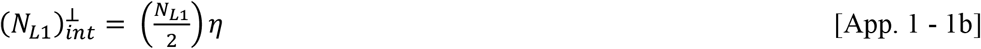

Under certain parameter regimes (such as those reflecting severe contact inhibition i.e. γ → 0) this different mode of contact inhibition will display different growth patterns to those considered in the main text. To highlight these differences some of the growth markers used in Figs. 4 and 5 are applied to simulations made using Eqn. 5 and 11 based on treatment of the surface layer as not causing contact inhibition to vertical growth of the cells resting upon it (Eqn. App. 1 a and b) (**App. Fig. 1b-e**).

**App. 1 – Figure 1:**
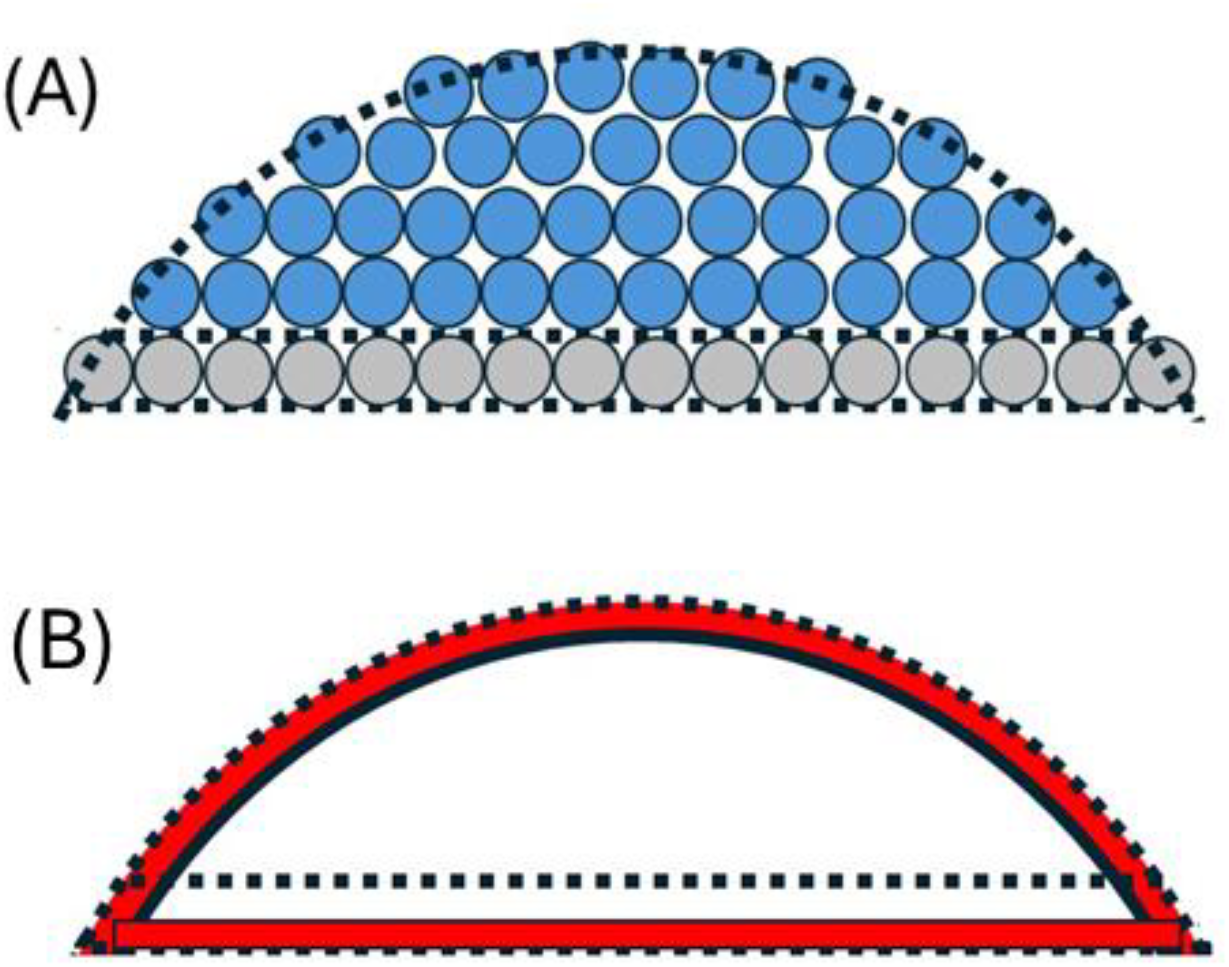
Sub-volumes of the spherical cap **(A) Total spherical cap volume, V**_**COL**_, **consists of the basal layer volume, V**_**L1**_, **and an upper cap volume, V**_**CAP**_: Equations defining the relations between V_COL_, V_L1_ and V_CAP_ are given by Eqn. 7 and 8. (**B) Schematic description of internal and perimeter regions of two spherical cap sub-volumes:** Each sub-volume, V_L1_ and V_CAP_, consists of internal (white) and perimeter (red) regions with the number of cells in each region given by Eqn. 5c and d together with Eqn. App1-1a and b (basal layer) and Eqn. 9a and d (upper cap).

## Footnotes

1 Such as would occur for cells in a well-stirred large liquid volume **[Hartwell and Unger, 1977; Ritacco et al. 2018]**.

2 As might be conferred by the nearby presence of a directing object – such as a surface or a food source.

3 Other smooth regular shapes would include an ellipsoidal cap and a two-dimensional Gaussian function.

4 In Appendix 1 we consider the effects of an alternative situation for which the plate surface does not act as a contact inhibitor of cell growth in the vertical direction and therefore the bottom facing aspect of the cells in contact with the surface will always constitute a perimeter region.

5 The importance of Ω is that it provides easy reckoning of colony shape i.e. if Ω = 2 the colony presents as a sphere on the surface, Ω = 1 indicates an exact hemisphere, and Ω < 1 indicates a minor spherical cap section of a greater sphere.

6 Defined in this manner Γ= 1 for spatially isotropic growth/division, Γ< 1 for vertically favored growth, and Γ > 1 for horizontally favored growth.

7 In general, there are three types of cell culture (i) liquid culture (cells suspended in solution), solid culture (cells grown on a surface) and mixed liquid/solid culture (cells are grown at a liquid/solid interface or within a hydrated scaffold) **[Adams, 1990; Bonifacino et al. 2004]**. Cell culture experiments involving a solid support are frequently classified as being either two-dimensional (2D), in which the solid support constitutes a flat surface upon which the cells grow, or three-dimensional (3D), where the solid support consists of a sufficiently soft and porous gel, inside which the cells may grow in a manner partially dictated by the shape of the scaffold **[Imamura et al. 2015; Kapalczynska et al. 2016]**. The present work is concerned with the 2D cell culture experiment i.e. cells growing on a flat surface.

8 Compare the typical weight force of a cell (buoyancy corrected ~10^−14^ N and non-buoyancy corrected ~ 10^−13^ N) vs. the random thermal force exerted by on an equivalent sized particle (in solution ~ 10^−12^ N and atmosphere ~ 10^−12^ N) at room temperature conditions. An understanding of this point is important especially for interpreting the results of particle-based simulation studies not featuring an intercellular attractive potential. In such cases newly formed cells will not necessarily ‘fall’ in place and colony geometry is not likely to be dictated balances of gravitational and friction effects of the types used to define the shape of structures such as sand dunes. However, for cases where a significant attractive potential exists (or is specified in a simulation) between cells and between cells and surface, the connected nature of the colony ‘gel’ will mean that the effects of gravity will become increasingly important in dictating the overall colony geometry as the colony becomes larger.

9 Greater detail on these junction types is given - occluding junctions (tight junctions and septate junctions), anchoring junctions (occur at either actin filament attachment sites e.g. adherens junctions and focal adhesions or intermediate filament attachment sites e.g. desmosomes and hemidesmosomes) and communicating junctions (e.g. gap junctions, chemical synapses and plasmodesmata in plants) **[Alberts et al. 2022]**.

10 With similar logic we might also suppose that cells within the basal surface layer would likely have a greater number of specific cell-surface linkages.

11 It is interesting to note that the geometrical relations developed in both cases **[Palumbo et al. 1977; Kamath and Bungay, 1988]** are only valid up to the point of a hemisphere and will produce an erroneous representation beyond this limit. Equation set 7 developed in the current paper solves this problem.

12 Following this definition, we will, for the sake of expediency, also include both particle dynamics type models, and tessellation-based mechanical models, for which the rules governing interactions can be determined/approximated via physics-based approaches.

13 Especially in the case of cell culture assays being monitored using high speed atomic force microscopy **[Yun et al. 2015; Guillaume et al. 2019; Puchkova et al. 2024; Hall and Foster, 2022; Hall, 2023b; Hall, 2023c]**.

